# Testing people’s aesthetic appreciation for biodiverse vegetation: Messiness is not a problem

**DOI:** 10.64898/2026.01.29.702517

**Authors:** Eva Breitschopf, Aaron Feicht, Eimear Tynan, Thomas Juel Clemmensen, Kari Anne Bråthen

**Affiliations:** UiT – The Arctic University of Norway, Department of Arctic and Marine Biology; UiT – The Arctic University of Norway, Academy of Arts; VERTE landskap-arkitektur AS

**Keywords:** urban vegetation, ecosystem functioning, species richness, native plants, niche complementarity, ecological filters framework

## Abstract

Biodiverse vegetation that supports high rates of ecosystem functions can inherently express a messy appearance due to high numbers of local native plant species and their spatial distribution connected to niche complementarity. This messiness is assumed to lower people’s appreciation for vegetation in urban contexts. Since such vegetation and a positive relationship between people and biodiversity could contribute to mitigating biodiversity loss, this assumed low public appreciation warrants investigation.
We designed and constructed biodiverse flowerbeds using only local native plants, and with the intention to enhance planting productivity, resistance and resilience. To investigate the influence of messiness, we created flowerbeds in four high levels of species richness (8,12,16,20) shown to be relevant for ecosystem functioning, and three levels of order (no, semi, full). We tested public appreciation for the flowerbeds using a self-guided, on-site survey.
We found a positive mean rating for all flowerbeds, but no effect of species richness on the ratings. Increased order, however, had a strong negative effect: The odds of a fully ordered flowerbed receiving a negative rating were 88% higher than of a flowerbed with no order. Increasing designed order was correlated with decreasing plant biomass in the flowerbeds.
These findings challenge the assumption that the appearance of biodiverse plantings is too messy for public appreciation in urban contexts. Specifically, we demonstrate that introducing order and reducing messiness can compromise aesthetic appreciation for biodiverse vegetation, potentially by compromising productivity as indicated by lower biomass production in ordered plantings.

**Synthesis and application:** Our study shows that biodiverse vegetation can be appreciated in urban contexts. Flowerbeds can effectively serve both people as ornamentation and biodiversity as habitat when they are designed based on ecological principles

**Research highlights:**

1. We designed and realized flowerbeds based on ecological principles.
2. All plantings received positive average ratings.
3. Increasing species richness in the flowerbeds did not affect participant’s aesthetic appreciation.
4. Increasing order in the design of the flowerbeds strongly lowered participant’s aesthetic appreciation.
5. Increasing order in the design was correlated to lower biomass productivity and more bare soil in the flowerbeds.

## Introduction

In the midst of the biodiversity crisis humanity is seeking solutions for biodiversity and people. With increasing urbanisation as one main factor threatening biodiversity through habitat degradation and loss (IPBES 2019) urban vegetation can be part of those solutions. Urban vegetation, including ornamental plantings, has the potential to support local biodiversity (Biella et al. 2025) and to strengthen people’s relationship with biodiversity (Ives et al. 2017). To realize this potential, urban vegetation needs to perform functionally as ecological plant communities (Klaus and Kiehl 2021) and aesthetically as part of people’s everyday life. In other words: plantings close to people need to be functioning ecosystems and their appearance needs to align with people’s preferences.

The challenge is that ecological functioning and an appealing appearance are often thought to be opposing features. Functional plant communities are thought of as messy and untidy, which collides with people’s assumed preferences and traditional design strategies favouring neat and ordered urban vegetation (Nassauer 1995; Li and Nassauer 2020). Since social and cultural considerations can act as constraints upon and preconditions for other imperatives (e.g. economic and environmental) (Boyer et al. 2016), the assumed lack of acceptance for plantings that are designed for ecological functioning can act as a constraint for their implementation. Hence, it can be expected that such plantings, especially for ornamentation, are only likely to be implemented and sustained if they are accepted by the public. Therefore, insights in how to design for ecological functioning and public acceptance in concordance are needed.

### Principles for designing for ecological functioning and biodiversity: Local native plants in communities of high species richness

The functioning of ecological systems encompasses many dimensions. *Ecosystem functions*, the ecological processes that transfer energy, nutrients and organic matter within and between ecosystems (e.g. production of biomass), result in outputs that contribute to people’s and other organisms’ lives in the form of *ecosystem services*/*nature’s contributions to people* (e.g. food, erosion control) (Behm 2020). To capture all these dimensions, we refer to *ecological functioning* as the property of an ecosystem that sustains and is sustained by its organisms and the ecological processes that they are part of (Jax 2005; Naeem and Wright 2003).

The ever-growing body of research in the field of biodiversity and ecosystem functioning (BEF) shows that ecosystem functioning is dependent on two primary components of species diversity: species composition (the identity of species) and species richness (the number of species) (Cardinale et al. 2011). The identity of species matters in terms of their fitness for the local environment and in terms of their functional roles in the community (Cadotte 2017). Which species naturally assemble to functioning plant communities is conceptualized by plant community theory considering species’ adaptations and local environmental conditions (Lortie et al. 2004). Species richness has been shown to increase many ecosystem functions (Weisser et al. 2017). One major explanation for increased ecosystem functioning in species-rich plant communities is niche complementarity (Loreau et al. 2022). Species occupy different ecological niches, such as e.g. time or depth of nutrient uptake (McKane et al. 2002) and fulfil different functions. Consequently, available resources are used complementarily and more completely when more species are present in a community and this effect increases over time (Amyntas et al. 2023).

Building on this foundational understanding of BEF, plant ecological theory and evidence suggest three main principles to design for ecological functioning: plant selections that are **1.)** based on **plant community theory**, **2.)** comprised of **local native plants**, in **3.) high species richness**.

**1) Plant community theory:** Designing ornamental plantings as functioning plant communities entails selecting species and placing the individuals in such a way, that interactions between them sustain ecosystem functions and the community’s persistence over time. This requires selecting species according to local environmental conditions and the interactions between the plant individuals. Plant community theory (i.e. integrated community theory (ICT), (Lortie et al. 2004)) can guide this selection process. The ICT describes the mechanisms of how plant communities assemble naturally, using the metaphor of filters: Factors in logically consecutive categories - 1.dispersal, 2. abiotic factors 3.biotic factors and 4. feedback mechanisms - filter the global species pool (i.e. all existing plant species) to a subset of species that live together in a plant community at the local scale. Applying the logic of these mechanisms to select plants for ornamental plantings is likely to result in a functioning plant community that is less dependent on human support. We employed this logic, the Ecological Filter Framework (EFF) (Breitschopf et al., in prep) as a tool in the design of our ornamental plantings and give a short description of the framework in Box 1.

**2) Local native plants** (plants of native species that stem from populations close to the site) are good candidates for ornamental plantings designed for ecological functioning because they are adapted to local environmental conditions, both physicochemical factors and other organisms. These adaptations are crucial for the plants’ ability to survive, grow and reproduce and can be to coarse-grained (e.g. photoperiod and temperature-regimes) as well as fine-grained environmental factors (e.g. biotic interactions and edaphic conditions) (Hamann et al. 2016). For example, Bjorkman *et al*. (2017), showed in an arctic common garden experiment that survival and reproduction rates of *Oxyra digyna* and *Papaver radicatum* individuals from local populations are higher than of individuals of the same species that originate from lower latitude populations, even with warming temperatures. Selecting plants from populations close to the designed site ensures they are adapted to coarse grain local environmental conditions and that they can exists in this area without human support. Further selection of species from this local native species pool, that fit the fine grain environmental conditions on site is likely to lower maintenance costs such as for e.g. winter protections, fertilizing or watering.

Furthermore, using local native plants in ornamental plantings can contribute to mitigating biodiversity loss. Firstly, designing with local native plants means providing habitat for them and other dependent organisms that might have been displaced by urbanisation (Biella et al. 2025). Particularly specialist organisms such as some local native pollinators are not supported by imported plant species (Burghardt et al. 2010). Secondly, it can avoid homogenisation among urban species pools and loss of species in adjacent ecosystems that are caused by the ubiquitous use and spreading of some very generalist ornamental species (van Kleunen et al. 2018; Beaury et al. 2021; Vellend et al. 2013).

We refer to *local native* plants to emphasize their co-evolution with and adaptation to local abiotic and biotic conditions underscoring their ecological fitness and roles. This emphasis is explicitly not intended to convey value-laden meanings such as belonging versus alien implying xenophobia, or good versus bad biodiversity, as discussed in e.g. Warren (2023).

**3) High plant species richness** in ecological communities has been shown to enhance ecosystem functioning (Weisser et al. 2017). Evidence and synopsis of hundreds of studies clearly demonstrates the positive effect of species richness and evenness on inter alia productivity, resilience and resistance to disturbances, invasive species and sensitivity to pathogens (Cardinale et al. 2011; Naeem et al. 1999; Weisser et al. 2017). All of which are desirable characteristics for ornamental plantings since they are likely to reduce maintenance costs and to increase the appearance of lushness. Moreover, these effects grow stronger over time (Reich et al. 2012; Weisser et al. 2017) and are likely to translate to ecosystem services/nature’s contributions to people, e.g. protection against erosion, water retention, carbon sequestration and more (Cardinale et al. 2012).

Designing for niche complementarity with a high number of species from differing functional groups and placement of individuals that allows for complementarity effects to occur can therefore lower maintenance costs by optimizing productivity and increasing resistance.

#### Box 1: The Ecological Filters Framework (Figure and shortened description from Breitschopf et al. (in prep))

The EFF is intended as a design tool to support decision-making in landscape architecture. It is inspired by the integrated community theory (ICT) (Lortie et al. 2004) that uses the metaphor of filters “filtering” the global species pool. Only species that “pass” these filters can be part of the local community. Consulting these filters for species selection or site manipulation can support ecological functioning when designing plantings.

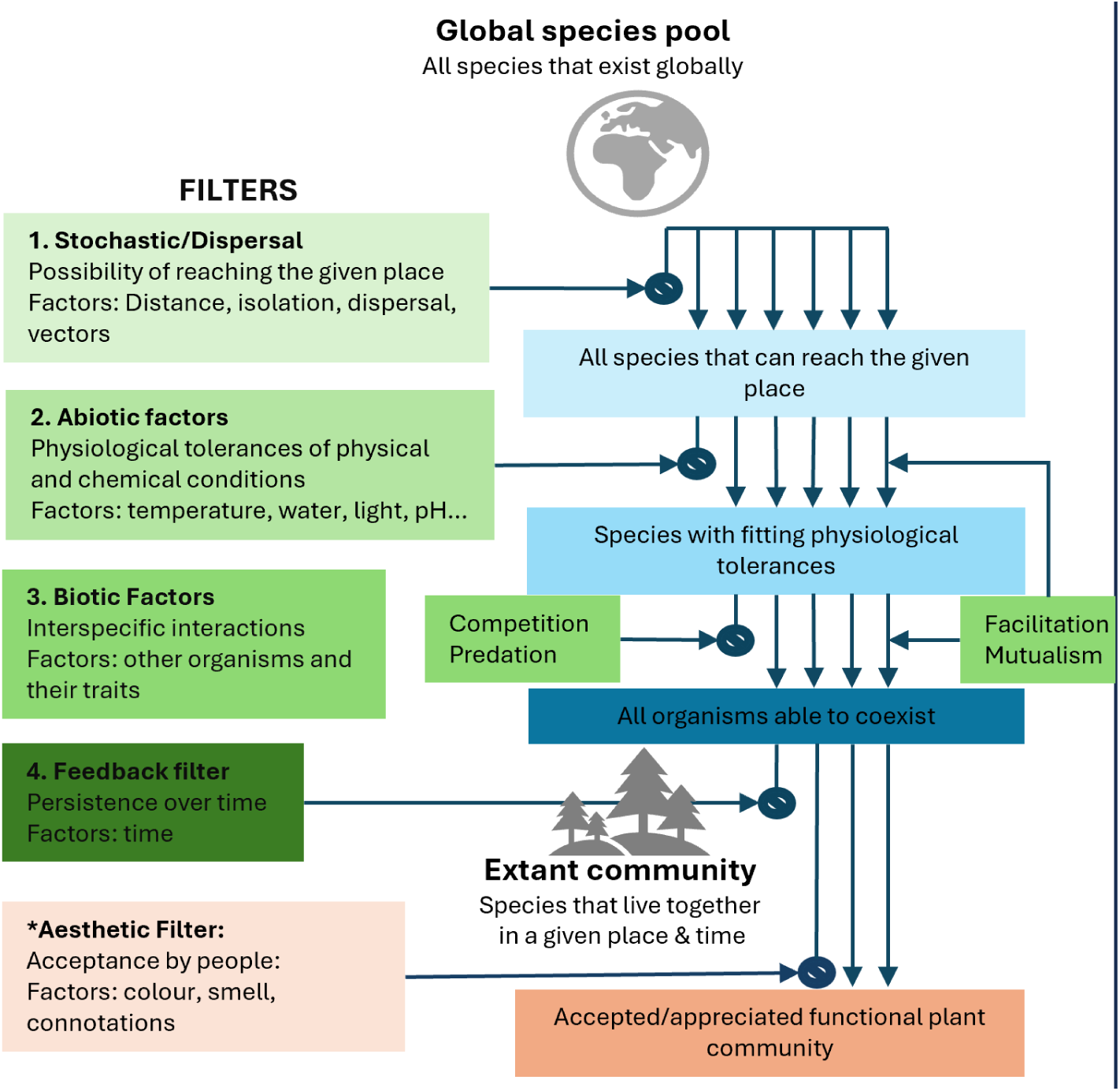

##### 1. Dispersal filter and stochasticity

Conditions that hinder a species from reaching the location: E.g., a large distance from the habitat of origin exclude species that cannot reach the location. This filter can artificially be overcome by importing species. It should be considered, however, if this is beneficial and necessary, since dispersal from the planting after import can impose a threat to local ecosystems. Local native species have passed this filter and do generally not pose an invasion risk.

##### 2. Abiotic filter

Not all species that reach a location can tolerate local physicochemical conditions (light, temperature etc.). Determining the abiotic conditions on site can aid in selecting species with matching niches. Local native species are adapted to local coarse grain conditions and therefore good candidates to further select from for fine grain abiotic conditions (e.g. pH).

##### 3. Biotic filter

Species that can reach a location and tolerate the local physicochemical conditions must be able to coexist with other organisms on site. Negative interactions with other species, such as competition for limiting resources, can lead to competitive exclusion. Positive interactions such as facilitation can be necessary for species to persist in the community. Supporting positive interactions and avoiding negative interactions can be steered by plant selection and designing for niche complementarity. E.g. including legumes is likely to facilitate productivity due to nitrogen fixation; excluding allelopathic and otherwise highly competitive species is likely to ensure survival of less competitive species. Designing for niche complementarity with many species that use resources differently can help avoid competition, increase the likelihood for facilitation to occur and thereby enhance ecosystem functioning (Godoy et al. 2020).

##### 4. Feedback filter

Communities change over time. The feedback filter therefore can include species that have been excluded (e.g. more time for dispersal (filter 1)) or exclude species that have been part of the community (e.g. accumulation of soil pathogens (filter 3)). Processes such as plant-soil feedback and legacy effects are especially relevant here (van der Putten et al. 2013). Foreseeing such feedback mechanisms can help maintain the intended plant community or anticipate and design its change.

##### 5. Aesthetic filter

Additionally, ornamental plantings are shaped by people’s aesthetic preferences. They need to be accepted by the public. Selecting species and design principles that cater to those preferences increases the likelihood of positive reactions to functional communities. E.g. large and colourful flowers increases peoples’ liking for a plant (Hůla and Flegr 2016) and cues to care such as an ordered appearance are thought to enhance acceptance (Nassauer 1995).

**In summary,** designing ornamental plantings as species-rich communities of local native plants offers multiple benefits. For the sake of easier reading, we will henceforth refer to this approach as “based on ecological principles.”

Plantings based on ecological principles will appear more similar to wild plant communities which are assumed to receive lower acceptance by the public. Planting design only started a development towards more “naturalistic” plantings in the last decades (Dunnett and Hitchmough 2004). More common and conservative planting designs typically rely on exotic species (Hunt 2000) and tend to avoid both local native plants and high species richness (Li and Nassauer 2020). It is therefore necessary to investigate whether the assumed lack of acceptance holds true and to develop strategies to design plantings based on ecological principles, that also are accepted and aesthetically appreciated by the public.

### Mediating messiness for public acceptance of ecologically functioning plantings

There are several reasons to suspect that planting design based on ecological principles may not align with people’s preferences since their above-described characteristics can infer messiness:

**1)** the appearance and cultural connotations of **local native plants**, and **2)** the complexity and lack of order resulting from **high species richness**.

**1) Local native plants** receive increasing attention but are still less likely to be incorporated in ornamental plantings than exotic ornamental plants due to lower availability for purchase, aesthetic preferences for exotics and cultural connotations against local native plants (Tartaglia and Aronson 2024). Peoples’ aesthetic preferences gravitate towards plant traits such as large, colorful flowers with radial symmetry (Hůla and Flegr 2016), uniform growth patterns, and a strong visual impact (Kendal et al. 2012) - traits that ornamental plants are specifically selected and bred for. Furthermore, exotic plant species carry the notion of the extraordinary (Hunt 2000), whereas local native plants, because they are present in local ecosystems, can seem ordinary due to people’s familiarity with them. Additionally, cultural perceptions might reinforce the avoidance of local native plants. Since they exist in independence of human care, their presence can be interpreted as neglect or poor maintenance. Since they are usually associated with wild, untended areas they might infer an appearance of neglect or even abandonment (Nassauer et al. 2009). Local native plants in managed land are labeled “weeds”, a term that carries negative connotations, implying something that is unwanted or out of place. Hence, both their aesthetics and their connotations make local native plants presumably less desirable for many people in designed spaces. This leads to low demand for their commercial availability, which feedbacks into the difficulty for including local native plants into ornamental plantings (Tartaglia and Aronson 2024).

**2) High species richness**, too, stands in contrast to widespread design traditions (Li and Nassauer 2020), even though an increasing number of studies finds a positive effect of biodiversity, especially species richness and evenness, on appreciation for a site or vegetation (Breitschopf & Bråthen, 2023; Southon et al., 2018; Hoyle et al., 2018, 2017a; Graves et al., 2017; Lindemann-Matthies et al., 2010). However, high diversity of plant species might introduce a visual complexity that diverts from the assumed desired clear and readable design language. A mix of varying heights, colors, textures, and shapes may be perceived as lacking coherence or unity. Traditional landscape design, often favors a more controlled, uniform appearance, that is easier to interpret and deemed more visually pleasing (Li and Nassauer 2020). The aesthetics of monocultures communicate human intention and control that cannot be mistaken for untended or abandoned landscapes (Nassauer 1995).

Finally, social norms exert a strong influence against change in planting practices. Because plantings and gardens can operate as communication systems and might be interpreted as a display of one’s character, it is not only the designer’s or owner’s opinion that matters (Lynch 1971; Nassauer et al. 2009). Even if individual people would not mind or even appreciate the presence of local native plants in high diversity, the assumption that others misinterpret their appearance for neglect and by extension might misjudge the owner’s character, causes prioritizing of neatness and order over ecological functioning (Nassauer 1995).

Mediating the apparent messiness of plantings designed based on ecological principles by introducing the appearance of order might be the balance to strike when designing for ecological functioning and public acceptance.

### Aim of study

In this transdisciplinary case study, conducted in the subarctic North of Norway, we aimed to investigate public acceptance for local native species rich ornamental flowerbeds and whether designed order can influence public acceptance.

To experimentally test the influence of order we manipulated designed order in two ways: the number of species in a planting, and the spatial arrangement of individuals within a planting. With increasing species richness and designed order, we tested the following hypotheses:

1. Flowerbeds designed based on ecological principles receive low acceptance from the public.
2. Increasing species richness lowers public acceptance for flowerbeds designed based on ecological principles.
3. Increasing order in flowerbeds designed based on ecological principles enhances public acceptance.

## Material and Methods

In 2022, we designed and installed 12 flowerbeds using only local native plants. In 2023, we conducted an on-site survey using a self-guided questionnaire to gather public opinions on the flowerbeds.

### Study site

The flowerbeds were installed on campus at UiT– The Arctic University of Norway (Figure 3) at 69. 6856 N, 18.9729 E on the island of Tromsøya with a coastal subarctic climate.

### Local native plants

#### Seed sampling and plant production

In autumn 2021, we collected seeds of 52 local native forb and graminoid species as candidates for the flowerbeds. Seed collection was conducted at various locations within a 50 km radius of the campus to ensure genetic diversity and local adaptations. From each population we harvested seeds of no more than 10% of the individuals, never depleting all seeds of one individual, thereby maintaining viability and genetic diversity of the donor population post-harvest. From these seeds we produced plant individuals in a climate-controlled greenhouse. A detailed description of the plant production is available in the Appendix.

#### Species selection

Plant species that produced insufficient numbers of individuals were excluded as candidates for the flowerbeds. We assessed the flowerbed’s abiotic conditions and selected species using an information matrix containing the plant species’ traits and characteristics relevant to the EFF, informed by Tyler et al., (2021), the local floras (Mossberg 2018; J. Lid 2005) and the authors’ experience.

We considered time of immigration, residence and invasive potential (dispersal – filter 1); light demand, moisture demand, soil reaction (pH), nitrogen demand, phosphorus demand, salinity tolerance (abiotic – filter 2); longevity, plant height, allelopathy, nitrogen fixation (biotic – filter 3); soil disturbance reaction, grazing or mowing dependency/tolerance (feedback – filter 4), flower timing, flower colour, flower/inflorescence size, and overall visual impact (aesthetic filter). Plants that were imported or with their niche outside the range of the flowerbed’s abiotic conditions and species with disadvantageous biotic interactions were excluded. Species with desirable aesthetic traits were prioritized.

### Study design

The flowerbeds were designed in four sets of three (Figure 1). The sets differed in species richness (8,12,16,20) in ranges that were shown to be relevant for elevated ecosystem functioning (Reich et al. 2012). Within a set, the flowerbeds differed in their level of order (no order, semi-order, and full order). The levels of order were realized by differing planting patterns (Figure 2).

**Figure 1:**
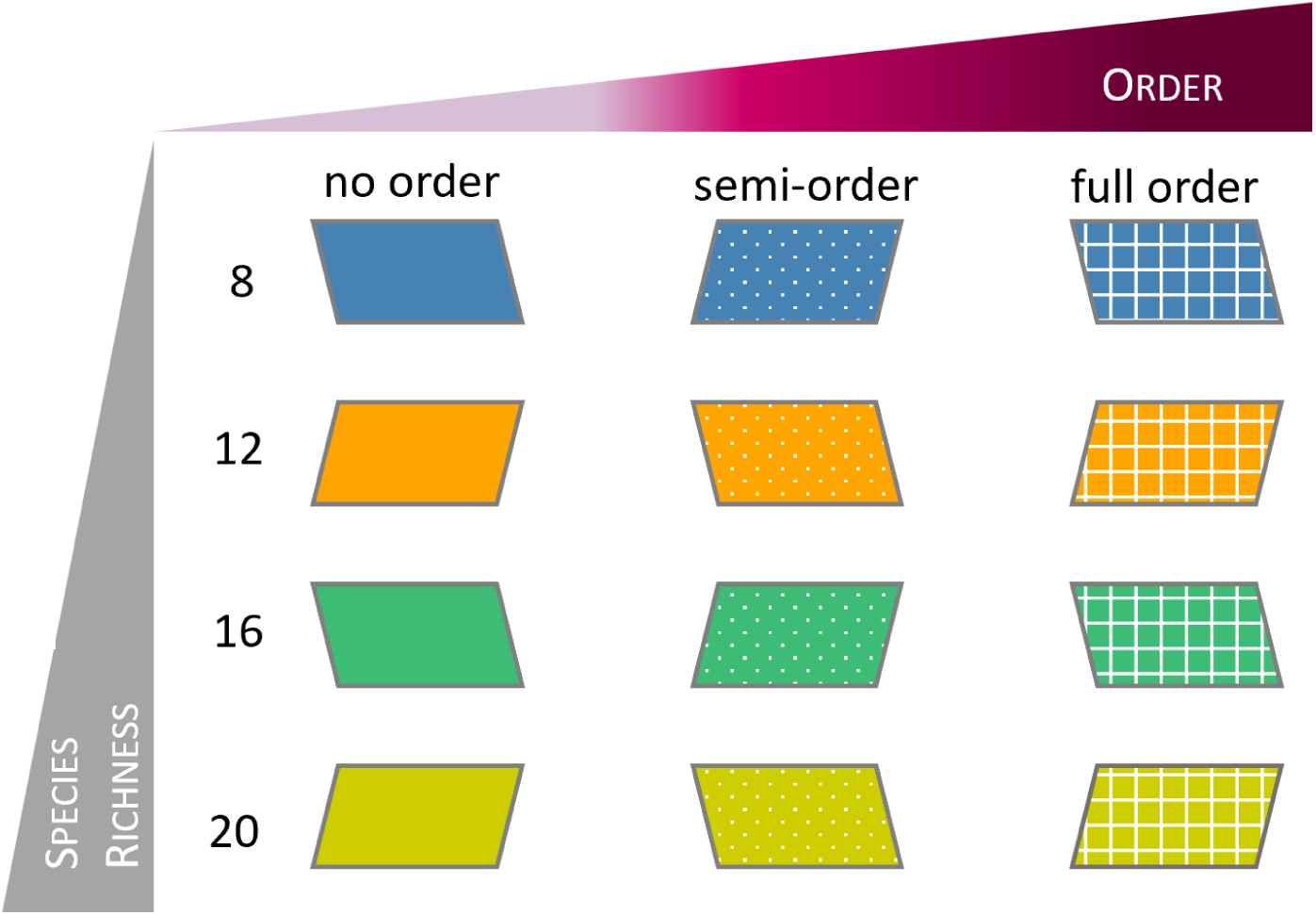
Diagrammatic visualization of the study design: Four sets with differing species richness (indicated by colours) containing three flowerbeds in increasing levels of order (indicated by pattern).

**Figure 2:**
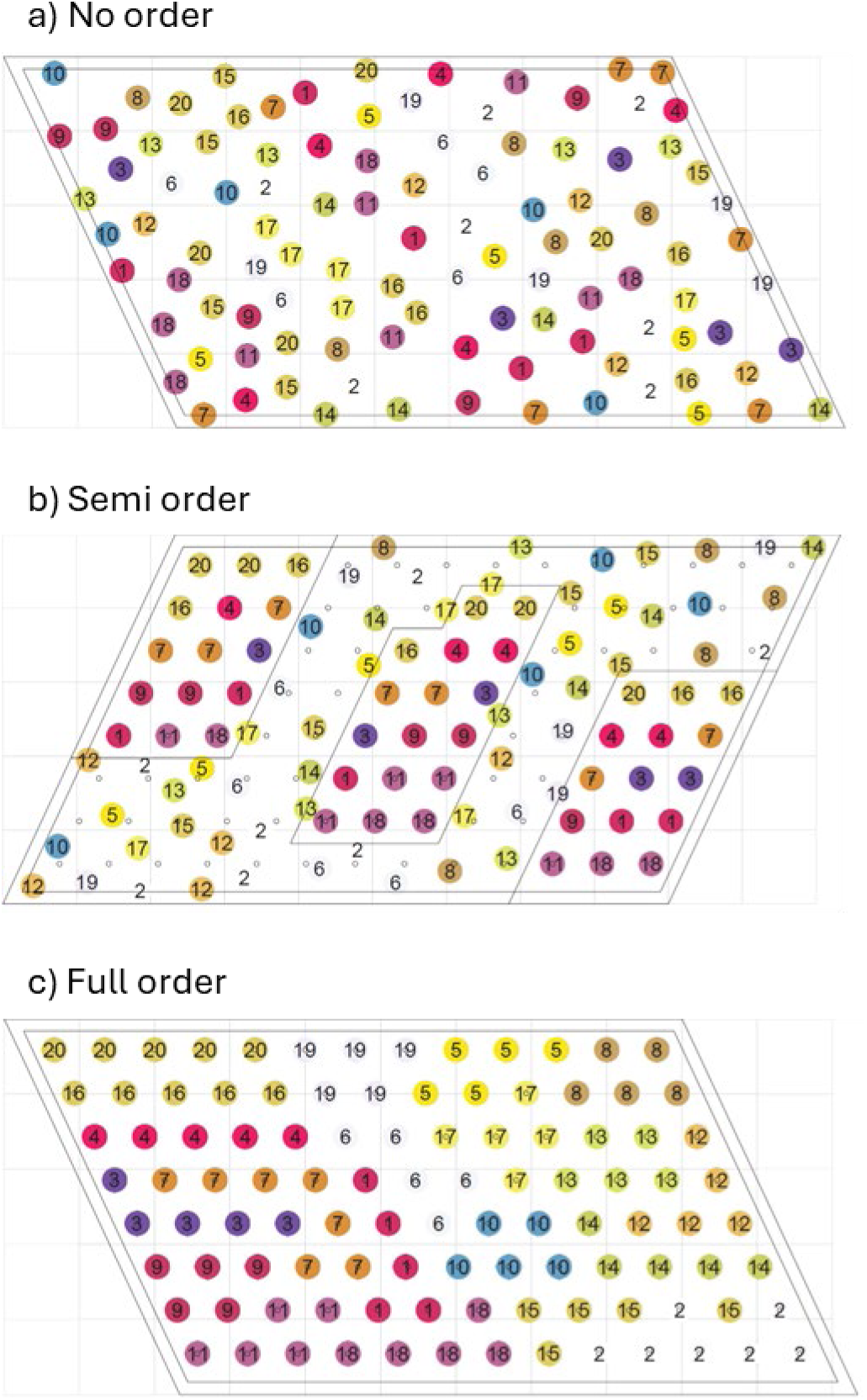
Planting plans for the 20 species set: a) flowerbed 4A, no order; b) flowerbed 4B, semi order; c) flowerbed 4C, no order. Dots represent individuals. Colour and number indicate species identity.

### Planting design

#### Constant factors

To avoid bias from factors other than species richness or order, the flowerbeds were standardized as follows: Each flowerbed consisted of 104 individuals (26 per m²) in equal species abundances. Flower colour identity (white, red, yellow, and green) and abundance were kept as constant as possible across all flowerbeds. We increased the number of species by consecutively adding four new species. A ratio of 1:4 grass to forb species was maintained to keep functional diversity constant. For every eighth species we included one legume species for potential nitrogen fixation. The list of the planted species is available in Appendix (Table S1).

#### Designing levels of order

The levels of order were realized by the spatial arrangement of the plant individuals within the flowerbed. This was designed and visualized with Grasshopper in Rhinoceros 3D (McNeel 2010).

To enhance the possibility for niche complementarity to occur, all individuals were placed to have at least two non-conspecific neighbours. Only at edges there could be less.

For **no order**, the plant individuals were distributed without any pattern, their positions were assigned randomly, observing the rule of non-conspecific neighbours.

For **semi order**, half of the plant individuals (red-flowered species accompanied by grasses) were placed on a grid while the other half (yellow, white, green) were placed without pattern.

For **full order**, all individuals were placed on a grid and ordered by height (tallest in the back) and their colour. To further maximize the appearance of order, a high visual impact species could not be next to species of low visual impact.

### Flowerbed design

#### Flowerbed placement

The flowerbeds were placed along a pedestrian/bike path, parallel to the biology building on campus (Figure 3). This location was chosen due to the high number of passersby in a divers demographic: students and staff working on campus; people commuting to other places of work or on a stroll, which includes people from the University Hospital and a kindergarten nearby and dog-walkers.

**Figure 3:**
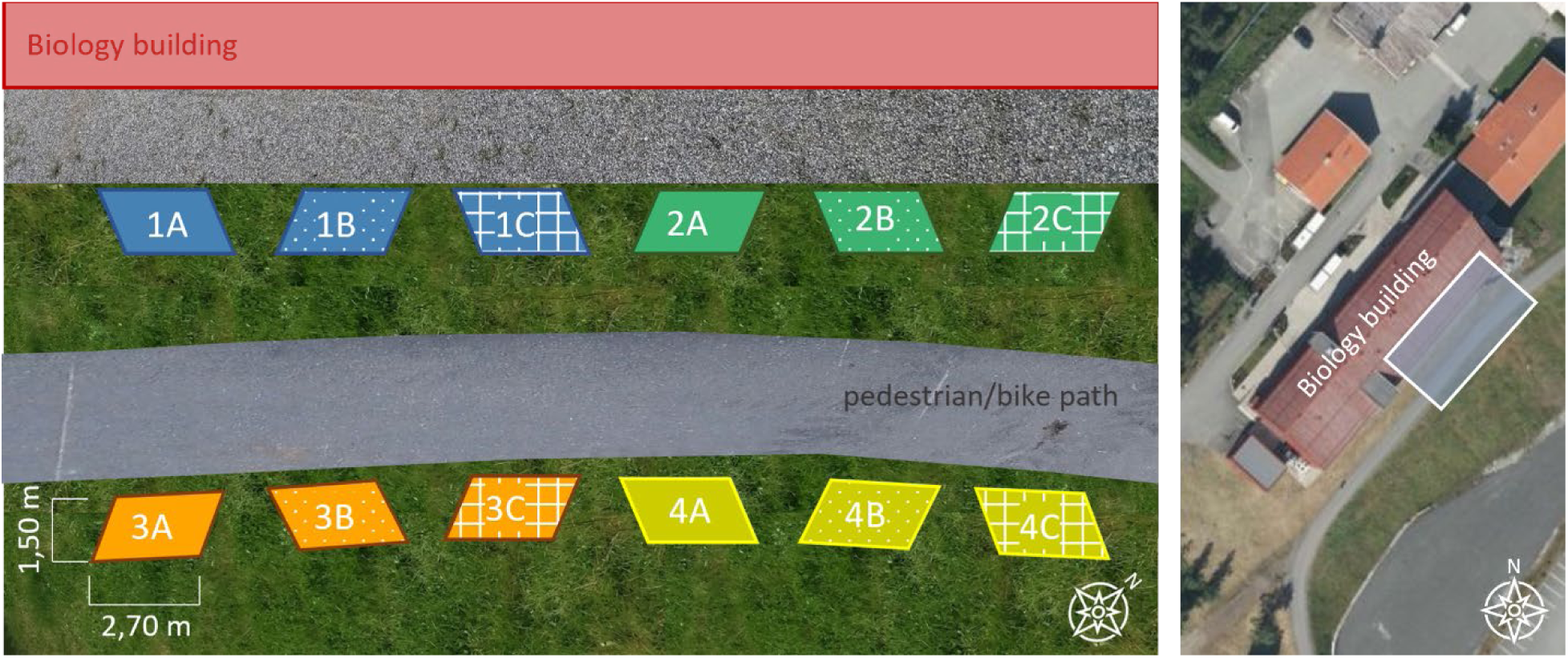
Placement plans, detailed (left) and overview (right, based on www.norgeskart.no).

Flowerbeds within a set were placed next to each other with 1m distance. The distance between sets was 2m (Figure 3).

#### Shape

The flowerbeds were designed as parallelograms (2,7 m x 1,5m, 65°) to maximize the visual size when walking by. A longer shape was chosen to complement the linear configuration of the location.

#### Planting the flowerbeds

The flowerbeds were excavated to a depth of 40 cm and filled with locally sourced compost soil due to the poor quality of the original soil on site. End of June 2022, the plants were transported to campus in pots and planted according to the planting designs.

### Vegetation Analysis

Mid-July 2023, we visually estimated the percent cover of bare soil and plant biomass in the flowerbed. End of September 2023, we cut down all standing biomass, dried it at room temperature for 20 days and weighed the dry mass.

### Survey

We initiated the survey in the end of May 2023 with setting up information signs about the study. The signs included QR codes linking to the questionnaire, available both in English and Norwegian, for participants to complete online, using their phones. We offered an identical paper version. Participants were not informed about the study design or its design principles, nor the type of plants that were used.

#### Questionnaire

The online questionnaire was created with nettskjema.no, developed and hosted by the University of Oslo.

The questionnaire started with a short introduction. Firstly, we asked for three word-associations for the flowerbeds in general. Then the participants were guided through the flowerbed sets and asked to rate each individual flowerbed on a 5-point Likert scale (1= I don’t like it at all, 3=Neutral, 5= I like it very much). Participants were requested to give a short reason for each rating. After rating all flowerbeds, participants were asked to state their favourite flowerbed and their favourite set. We asked three more questions on the flowerbeds in general, inquiring whether the participants would like to see more of this kind of plantings in town/on campus/in their own garden. Then, we included questions about the participant’s personal background (age, citizen of Norway/Troms og Finnmark/Tromsø, familiarity with Tromsø/the used plants, work/study on campus, experience with design of outdoor space/gardening/ecology). Following this main part of the questionnaire, we informed the participants about the ecological design principles and reasoning behind the flowerbed designs and asked whether that information would have changed their assessment. Lastly, we investigated participants’ prior knowledge of the study. The questionnaire is available in Appendix (Table S2).

### Research ethics

We adhered to the guidelines set forth by the Norwegian National Committees for Research Ethics. The questionnaire was assessed and approved for data security by SIKT, the Norwegian Agency for Shared Services in Education and Research.

#### Consent procedure

Participants were informed about the intent and origin of the study, the nature of the questionnaire, as well as their anonymity and voluntary participation, on the invitation signs and on the front page of the questionnaire. After the final question, participants were reminded that participation is voluntary, anonymous and that with sending in their answers, they consent to their answers being used in this study. Participants had the option to withdraw from the questionnaire at any time before submitting online or delivering the paper version, ensuring no registration of their answers.

### Statistical analysis

All statistical analysis was performed in R 4.4.3 (R Core Team 2025).

We used a linear model to visualize the effects of species richness and order on the average ratings. To estimate the effect sizes of species richness and order on ratings we employed ordinal logistic regression using the “VGAM” package (Yee 2010). Since the proportional odds assumption was violated, that is the odds ratios across all categories were not consistent, we applied a generalized ordinal logistic regression (Williams 2016). For easier interpretation we collapsed categories: negative rating= 1 (“I don’t like it at all”) + 2 (“Do not like); neutral rating = 3 (“neutral”); positive rating = 4 (“I like it) + 5 (“I like it very much”).

## Results

### Participants

A total of 177 people participated in the survey. Most participants where from Norway (58%, n=102,) and described themselves as familiar with Tromsø (familiar: n=53, very familiar: n=45, somewhat familiar: n=38, not familiar: n= 14, not at all familiar = 18).

Participants’ age ranged from 11-20 to 71-80 years, with most in the age group 21-30 (43%, n=77). More participants reported working off-campus (49%, n= 87) than on-campus (45%, n=79).

Most participants rated their experience in design (n=134; 75,71%); gardening (n= 149; 84,18%), or ecology (n=71,18%) to be at or lower than hobby level or did not specify. Only 19 participants rated themselves at professional level for design (above hobby level: n= 43), 7 for gardening (above hobby level: n=28), and 5 for ecology (above hobby level: n=51).

Few participants considered themselves unfamiliar with the plants (not at all familiar: n=1, not familiar: n=7, somewhat familiar: n= 44, familiar: n= 71, very familiar: n=43).

### Vegetation

The percentage of bare soil was significantly correlated with the harvested biomass (r= -0.7474; CI_95_: -0.9247, -0.3037). The produced biomass decreased strongly with full order and only slightly with increasing species richness (Table 1, Figure 4). Further findings on the plantings functional performance and photographs of all plantings (corresponding to Figure 5) are available in the Appendix.

**Figure 4:**
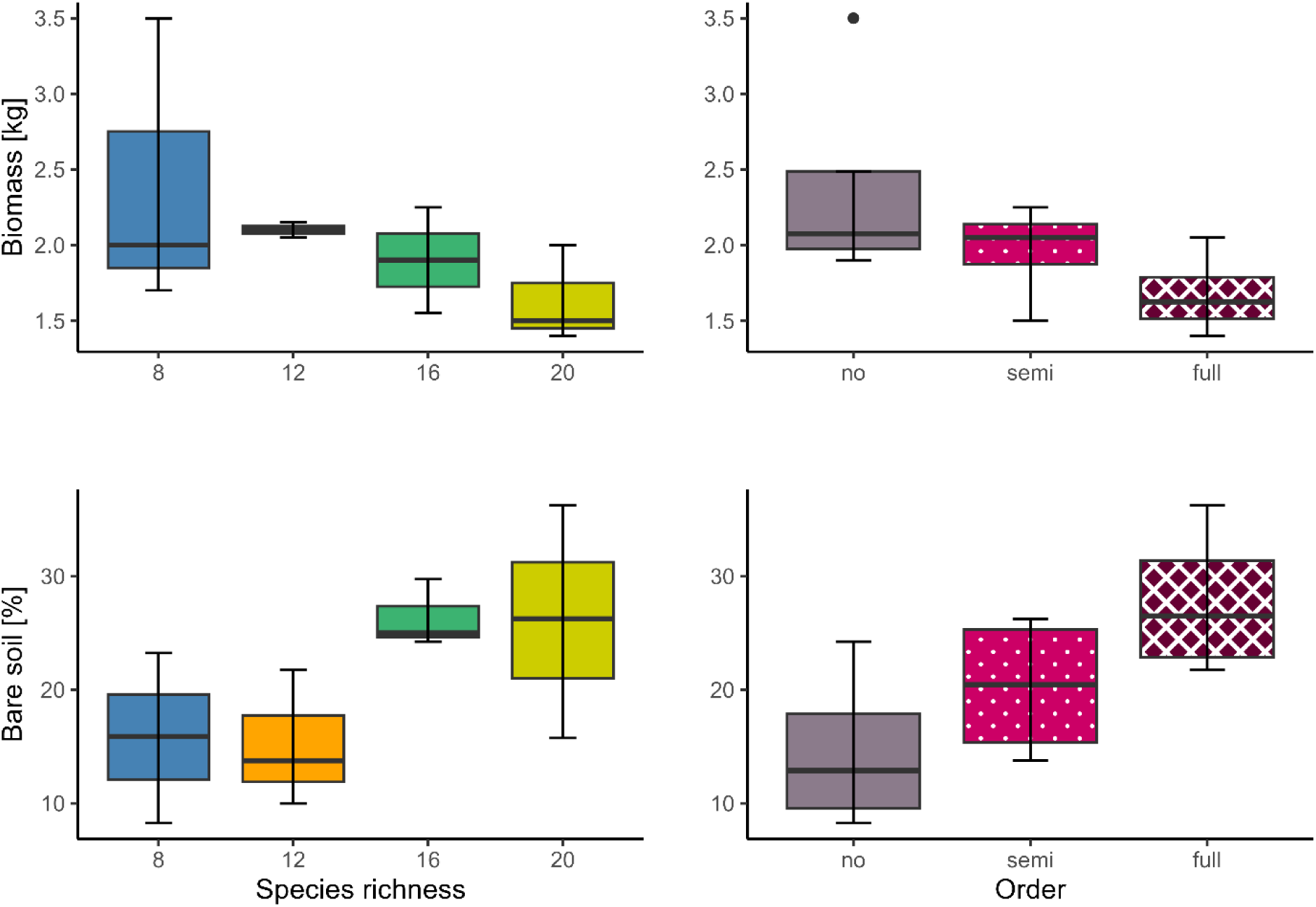
Boxplots of biomass and bare soil cover per species richness and order.

**Figure 5:**
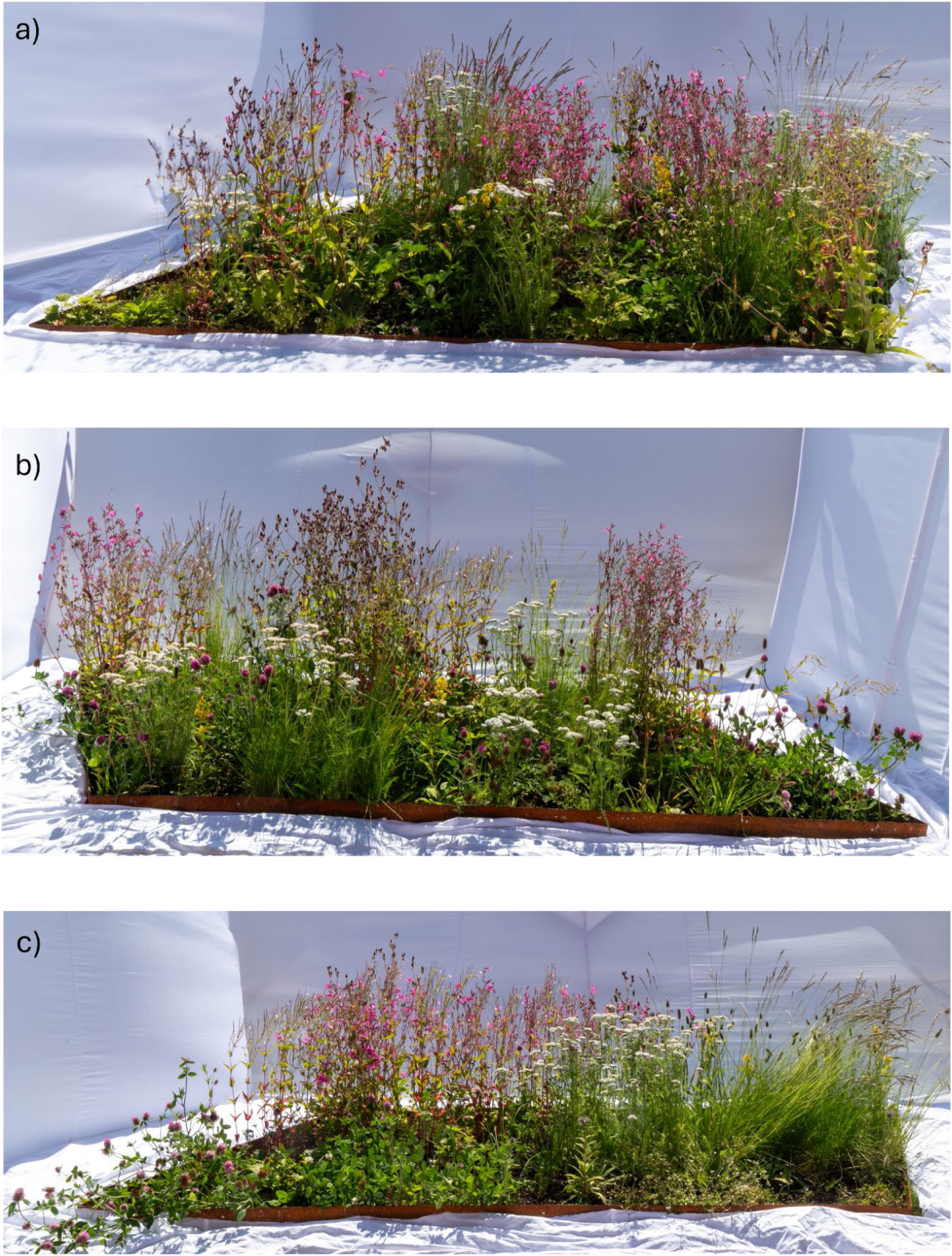
The flowerbeds of set B (12 species) with **a)** no order, **b)** semi order and **c)** full order, July 2023. Photographs of all flowerbeds are available in the appendix (Appendix). Photo credit: Oddleif Larsen.

**Table 1:** Model estimates for the linear model (effect of species richness and order on biomass) with confidence intervals at the 95% level (CI). CIs not overlapping zero indicate significant effects and are marked with * and bolded. R^2^=0.603.

| BIOMASS [KG] | ESTIMATE | CI 2,5% | CI 97,5% |
| --- | --- | --- | --- |
| **INTERCEPT\*** | 3.2625 | 2.3047 | 4.22026 |
| **SPECIES RICHNESS\*** | **-0.0625** | -0.1224 | -0.00264 |
| **ORDER SEMI** | -0.4250 | -1.0807 | 0.23073 |
| **ORDER FULL\*** | **-0.7125** | -1.3682 | -0.05677 |

### Participants’ opinion on the flowerbeds

All flowerbeds received positive average ratings, meaning an above neutral response. Unordered flowerbeds consistently received the highest average ratings, except for the 16-species set. Fully structured flowerbeds always received the lowest average rating (Figure 6). The average rating across all flowerbeds was 3.404.

**Figure 6:**
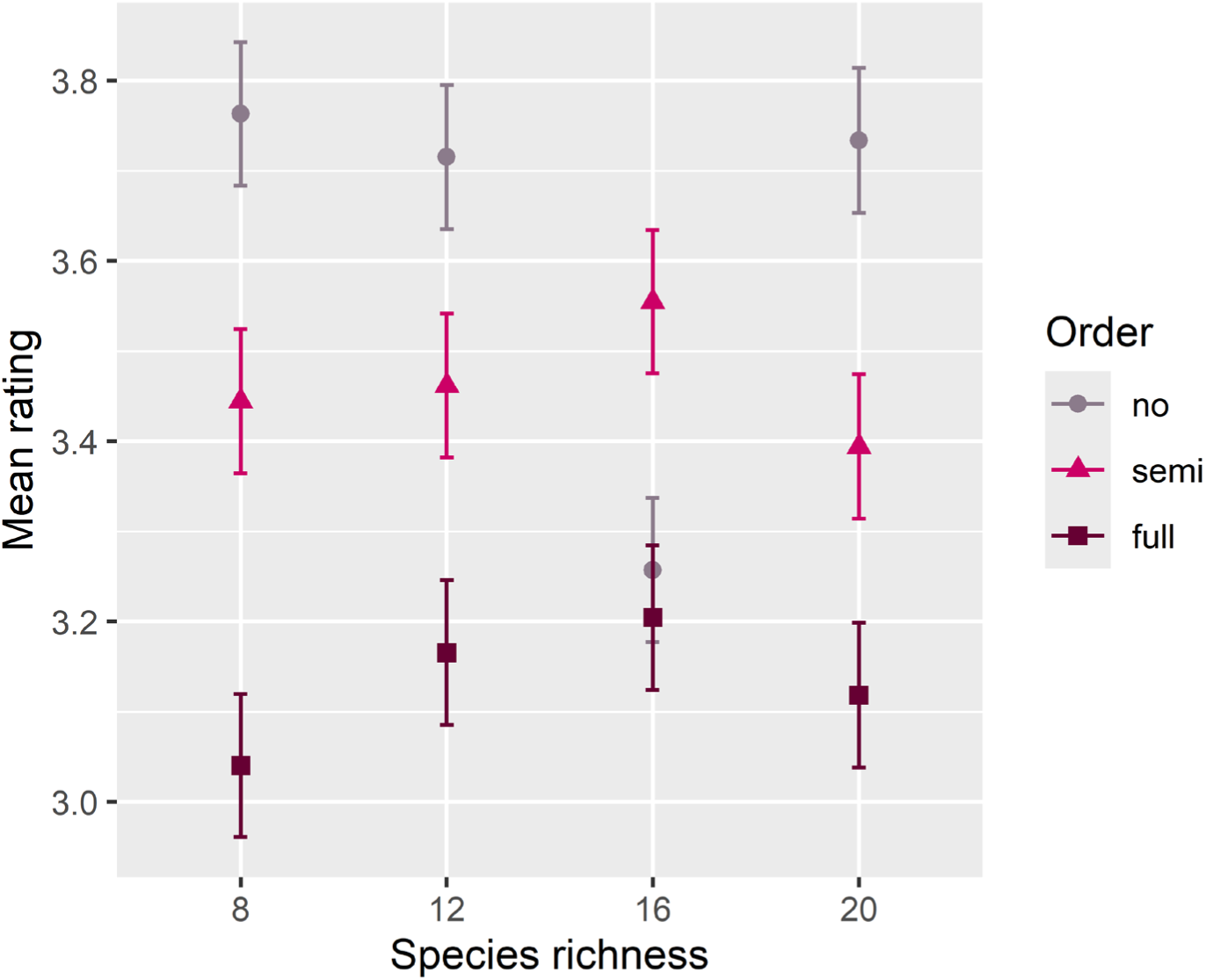
Average ratings (arithmetic mean ± standard error) for all plantings by species richness and order.

These results are independent of whether people with ecology experience above hobby level are included or excluded from the dataset (Appendix, Table S3, Figure S5). In the subsample excluding participants with ecology experience, the average rating across all flowerbeds was 3.364.

Our participants’ ratings for the flowerbeds were significantly influenced by the design of the flowerbed (Table 2). Increased order had a strong negative effect. The odds for a fully ordered flowerbed receiving a negative rating were 88% higher compared to a flowerbed with no order. We did not find an effect of species richness on the participants’ ratings (Table 2).

**Table 2:**
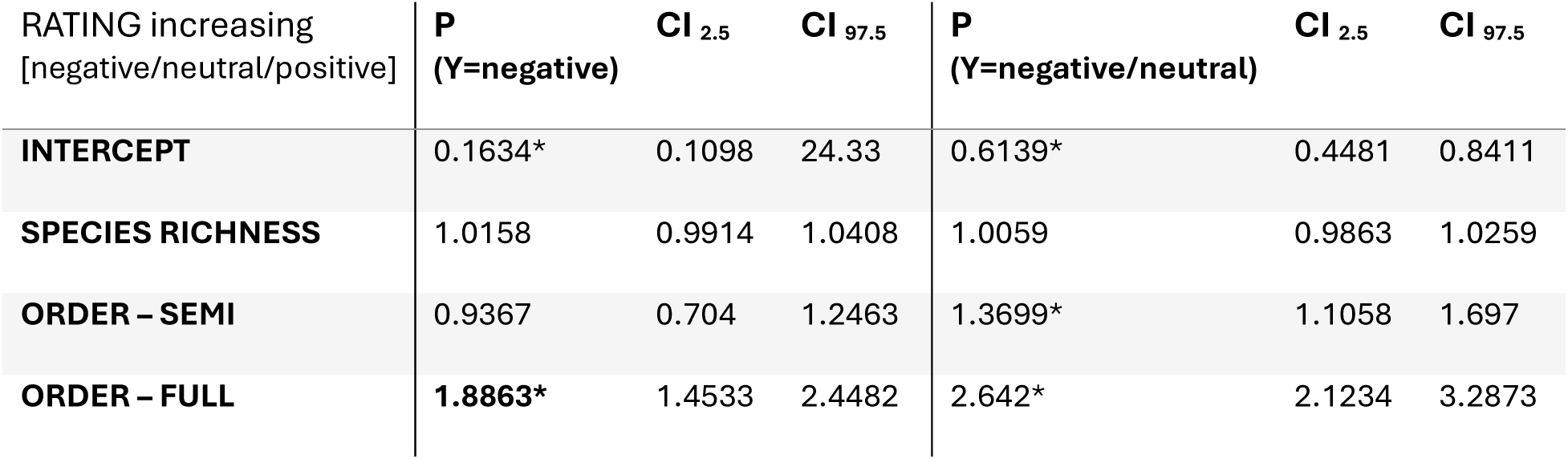
Model estimates for the generalized ordinal regression model (effect of species richness and order on ratings). The values given relate to an increasing order of ratings from negative to positive. Confidence intervals at the 95% level (CI). CIs not overlapping one indicate significant effects and are marked with *. AIC_Δ_ = -79 compared to null model.

Also, these results are independent of whether people with ecology experience above hobby level are included or excluded from the dataset. In the subset excluding participants with ecology experience, the odds for a fully ordered planting receiving a negative rating were 81% percent higher than for a planting with no order (Appendix, Table S4)

## Discussion

Our case study could not confirm the general assumptions on low public acceptance for local native, species rich plantings.

Contrary to hypothesis 1, all our flowerbeds received positive ratings. Additionally, we could not confirm hypothesis 2: we found no effect of increasing species richness on participants’ ratings for our flowerbeds. Finally, we found that increasing designed order had a strong negative effect on the ratings, which opposes hypothesis 3.

Thereby our results indicate that assumptions on public acceptance for plantings designed based on ecological principles do not always hold true and might be in the process of changing.

In the following we first discuss our findings in relation to the posed hypotheses and then elaborate on implications of our findings.

### I. Flowerbeds designed based on ecological principles DO NOT receive low acceptance by the public

Our hypothesis assumed that messiness inherent in functioning ecosystems contrasts with people’s aesthetic preferences leading us to expect that people would not like the flowerbeds we designed. This assumption was not confirmed. Rather, our flowerbeds were liked by the participants with all flowerbeds receiving average ratings above neutral.

We found few and mostly dated studies that empirically investigated public opinion on planting designs, and none specifically testing designs based on ecological principles to optimize ecological functioning. Aside from Nassauer’s studies from the 1990s and even earlier, we identified only two studies from the 2000s that compared “naturalistic” to „formal” design. Özgüner and Kendle (2006) and Özgüner et al. (2007) investigated the general public‘s and landscape professionals’ opinion on “naturalistic” vs. “formal” park design, finding no general preference for either style. However, pronounced differences existed among professionals: Those involved with conservation projects favoured naturalistic plantings, while professionals from local authorities and private companies were sceptical about public acceptance and positive experiences with naturalistic plantings. Minding the different focus (park views vs. flowerbeds, naturalistic vs. design based on ecological principles), these insights combined with our results, could suggest that the general assumption of public rejection of “messy ecosystems” might be upheld among professionals more than by people’s actual dislike. Furthermore, a more recent study empirically testing people’s preferences in connection to naturalness confirms that people find attractive, even though not tidy, what they perceive as natural and therefore as beneficial for insects (Hoyle et al. 2019). However, another study by Hoyle et al. (2017b), found that, while the majority of participants were positive towards non-native species, attractiveness and perceived nativeness were unrelated. This, in line with our results, indicates changing public attitudes from rejection to appreciation of natural-looking planting designs. Given the significant lack of research testing public attitude towards plantings designed based on ecological principles, these explanations remain speculative and call for a more thorough investigation and further research.

### II. NO EFFECT of increasing species richness on public acceptance for flowerbeds designed based on ecological principles

Contradicting our hypothesis, we found no effect of increasing species richness on participants’ liking. The complexity introduced by adding more species to the design does not appear to hamper public appreciation.

Previous studies across various contexts (wild (Graves et al. 2017), assembled (Lindemann-Matthies et al. 2010), artificial (Breitschopf and Bråthen 2023) and urban (Hoyle et al. 2017a; Qiu et al. 2013) plant communities) found negative (Qiu et al. 2013), neutral (Graves et al. 2017) and positive (Hoyle et al. 2017a; Breitschopf and Bråthen 2023) effects of species richness on people’s preferences. Given our case study’s strictly designed setting with the flowerbeds intended as ornamental vegetation, we expected the assumed preference for order to have a larger impact than in vegetation without urban context. However, our results align with findings from studies on wild plant communities, which show no impact of species richness alone. Furthermore, we tested the effect of species richness in levels that are high for ornamental plantings. Previous studies on the perception of biodiversity in plant communities indicate that differences in species richness may be less distinguishable in higher (>16) than lower numbers of species (Lindemann-Matthies et al. 2010; Breitschopf and Bråthen 2023). This low perception of differences in species richness could explain the lack of a species richness effect in our plantings.

### III. Increasing order in flowerbeds designed based on ecological principles DECREASES public acceptance

Our findings show that introducing order into flowerbeds designed based on ecological principles contrast with what people appreciate, suggesting that there can be too much orderliness for “messy ecosystems”. While the existent orderliness, such as neat edges and geometric shapes, may have sufficiently signaled human intention, the additionally introduced order within the plantings led to participants’ dislike. The neat appearance of the plants on a grid in the ordered plantings might clash with the “wild” appearance of local native plants, with the perceived control appearing forced and overly dominant, which might seem inappropriate for otherwise “wild” species. This interpretation could be further explored through the qualitative descriptions and word associations gathered in our survey.

Additionally, we found that order in the flowerbeds correlated with their productivity. Increasing order decreased the produced plant biomass and increased the bare soil area, which in turn could be responsible for the lower ratings. The ordered placing, despite our intention to maintain the possibility for niche complementarity, likely resulted in fewer non-conspecific individuals able to use resources complementarily, thereby impairing the plant community’s ecological functioning. Introducing order in that way, therefore, might lower plantings productivity and thereby decrease people’s liking for them.

### Implications

The findings of our study suggest that productivity, and consequently the appearance of lushness and fullness, may be a primary beneficial factor for people’s appreciation for ornamental plantings. This would imply that designing for ecological functioning is crucial. Compromising ecological functioning through increased order or low species richness would not only increase maintenance cost and effort but also compromise aesthetic performance.

With the absence of a negative effect of high species richness on public acceptance for ornamental plantings, concerns of dislike for species rich ornamental plantings appear to be of low relevance. Coupled with the positive ratings for our flowerbeds which contain only local native plants, we are confident to suggest exploring designs that rely on highly diverse and functioning plant communities. Such plantings can support biodiversity and benefit from high ecosystem functioning, while also being accepted by the public.

Aesthetic preferences, social norms and design traditions might be in the process of changing. Many of the discussed hindrances for public acceptance for ecologically functioning ornamental plantings are assumptions that we did not find explicitly tested or that might be outdated in the current time of increasing environmental awareness. Our results indicate that the public may be more accepting of ornamental plantings that are designed to support biodiversity and ecological functioning than generally assumed.

Furthermore, as attitudes appear to shift towards more naturalistic ornamental plantings, the construction of more plantings designed based on ecological principles, and thereby the more frequent exposure for people to them, could help to further reduce the cultural/social hindrances to their implementation. With plantings designed based on ecological principles becoming prominent, especially in public spaces, the message communicated by the plantings would no longer be “messiness” and “loss of control” but the care and appreciation for functioning ecosystems and biodiversity. This, in turn, could result in a closer connection of people and biodiversity in urban areas.

## Conclusions

In this case study in the climatically extreme environment of the Subarctic, we demonstrated that ornamental planting design based on ecological principles does not need to be held back by assumptions of their low aesthetic performance. Using high numbers of species and local native plants supports biodiversity and ecosystem functions such as productivity and resilience, which, in turn seem to enhance public aesthetic appreciation. Instead of the assumed dislike for the plantings’ inherent messiness our study demonstrated that our participants appreciate the appearance of functioning plant communities as ornamental vegetation, and that decreasing ecological functioning by introducing order might be the reason for participants’ decreased liking. With the implicated benefits for both functional and aesthetic performance we advocate design strategies that rely on knowledge found in the field of ecology. Tools like the EFF have proven instrumental in designing for ecological functioning and public acceptance.

Recognizing the lack of research, specifically testing design strategies for ameliorating the biodiversity crisis, we aim to work on further investigating the causal mechanisms for like or dislike for our plantings with the qualitative reasoning given by our participants.

## Acknowledgements

This research was funded by a PhD Scholarship at UiT – the Arctic University of Norway and supported by a business-mentorship program at UiT, funded by the county administration of Troms and Finnmark, enabling fruitful cross-sector collaboration. We appreciate the support from the Climate Laboratory at Holt, especially Leidulf Lund, who facilitated plant production. We thank Sophia Zielosko, Orhan Grignon, Marie-Pierre Bazan, Reviel Müller, Helmi Köykkä and Alen Perkovic for their work in the flowerbeds and Anita Veiseth for her valuable insights in the designing process.

## Author’s Contributions

All authors collaboratively conceived the ideas and designed methodology. EB and KAB established support for placing the flowerbeds at UiT campus. The flowerbeds were designed collaboratively, with ecology led by EB, supported by KAB; and aesthetics led by AF and ET, supported by TJC. The questionnaire was developed collaboratively, led by EB. EB collected the data and analysed the data, supported by KAB. EB led the manuscript writing. All authors contributed critically to the drafts and approved the final publication.

## Data availability statement

The supporting data sets and corresponding code for statistical analysis will be made available on DataverseNO (https://doi.org/10.18710/L3TOJT) upon peer-reviewed publication.

