## Appendix for "Testing people’s aesthetic appreciation for biodiverse vegetation: Messiness is not a problem"

### Producing local native plant individuals

The sampled seeds were air-dried at room temperature for 2 days, then cleaned from dirt and excess biomass. They were stored in paper bags at -20° C from September to November. When outside temperatures constantly fell below freezing, the seeds were stored dry and dark outside to expose them to winter temperatures and temperature fluctuations. To initiate germination, the seeds were further stratified in 0,5°C and darkness for 2 weeks in a wet mix of sand and vermiculite in paper cups with ventilation holes in the cover. The seed-sand-vermiculite mix was then spread onto 4 cm soil in 30x40 cm trays and covered with extra vermiculite to keep moisture levels constant and avoid algal and fungal growth. Under continuous light in 15-18° germination started within 3-10 days. When a seedling reached a sufficient size, it was transferred into a soil-perlite mixture (7:3) in 7x4x4 cm pot planting-trays, one seedling per pot. When the individuals outgrew these (within 2-4 weeks), they were transferred to 1-litre planting pots and stored for growth in a greenhouse with daylight at 12 (+-3) °C. They were watered, when necessary, 1-5 times a week.

***Table S2***: *List of species with general information*

| in Set | Nr of species | Species | Colour category | Functional group |
|---|---|---|---|---|
| A+B+C+D | 8 | *Trifolium pratense* | Red | forb, legume |
| A+B+C+D | 8 | *Cerastium fontanum* | White | forb |
| A+B+C+D | 8 | *Epilobium montanum* | Red | forb |
| A+B+C+D | 8 | *Silene dioica* | Red | forb |
| A+B+C+D | 8 | *Solidago virgaurea* | Yellow | forb |
| A+B+C+D | 8 | *Achillea millefolium* | White | forb |
| A+B+C+D | 8 | *Avenella flexuosa* | Green | graminoid |
| A+B+C+D | 8 | *Phleum alpinum* | Green | graminoid |
| B+C+D | 12 | *Geum rivale* | Red | forb |
| B+C+D | 12 | *Saussurea alpina* | Red | forb |
| B+C+D | 12 | *Allium schoenoprasum sibiricum* | Red | forb |
| B+C+D | 12 | *Festuca ovina* | Green | graminoid |
| B+D | 16 | *Ranunculus acris* | Yellow | forb |
| B+D | 16 | *Alchemilla subcrenata* | Yellow | forb |
| B+D | 16 | *Trifolium repens* | White | forb, legume |
| B+D | 16 | *Deschampsia cespitosa* | Green | graminoid |
| D | 20 | *Trollius europaeus* | Yellow | forb |
| D | 20 | *Campanula rotundifolia* | Red | forb |
| D | 20 | *Filipendula ulmaria* | Red | forb |
| D | 20 | *Leymus arenarius* | Green | graminoid |

*Table S2:* *Questionnaire summary - overview of sections and questions with possible answers*

| Nr. | Question | Answer type |
|---|---|---|
| I | Please write down the first 3 words that come to mind about the 12 plantings in general. | free-text |
| II<br>a)<br>b)<br><br><br>c)<br>d)<br><br><br>e)<br>f) | *Please go to the plantings in SET 1 (blue)*<br>How do you like planting 1A?<br>Could you give a reason for your rating or a word that describes planting 1A?<br><br>How do you like planting 1B?<br>Could you give a reason for your rating or a word that describes planting 1B?<br><br>How do you like planting 1C?<br>Could you give a reason for your rating or a word that describes planting 1C? | <br>Likert scale 1-5<br>free-text<br><br><br>Likert scale 1-5<br>free-text<br><br><br>Likert scale 1-5<br>free-text |
| | II a) to f) was repeated for set 2 to 4. | |
| III a)<br>b) | Which planting is your favourite?<br>Which set is your favourite? | multiple choice: 1A, 1B, 1C….4A, 4B, 4C<br>multiple choice: 1,2,3,4 |
| IV<br>a)<br><br>$b^{1}$)$c^{1}$)$d^{1}$)<br><br><br><br>$b^{2}$)$c^{2}$)$d^{2}$) | General questions about the plantings<br>Do these plantings meet your expectations of what a planting should look like?<br>Would you like to see plantings like these<br>a) IN TOWN<br>b) ON CAMPUS<br>c) In your OWN GARDEN?<br>Could you give a reason why or why not? | <br>multiple choice: Exceed, meet or fall short of my expectations<br><br><br>multiple choice: Yes, No, I don't know<br><br>free-text |
| V<br>a)<br>b)<br>c)<br>d)<br>e)<br><br>f)<br>g) | Questions about participants<br>How old are you?<br>Are you from Norway?<br>Are you from Troms/Finnmark?<br>How familiar are you with Tromsø?<br>How familiar are you with the plants that we used?<br>Do you work or study at on campus?<br>How would you rate your knowledge and experience with<br>- designing outdoor space?<br>- planting and gardening?<br>- plant ecology? | <br>free-text<br>multiple choice: yes, no, no answer<br>multiple choice: yes, no, no answer<br>Likert scale 1-5<br>Likert scale 1-5<br><br>multiple choice: yes, no, no answer<br><br><br>Likert scale 1-5<br>Likert scale 1-5<br>Likert scale 1-5 |
| VI<br><br>a)<br><br>b)<br>c) | *Information text on the ecological and sustainability context of the planting design*<br>Do you think this information would have changed how you assessed the plantings?<br>Do you have prior knowledge of this study?<br>Have you answered this questionnaire before? | <br><br>multiple choice: better, the same, less<br><br>multiple choice: yes, no<br>multiple choice: no, once, twice, >twice |

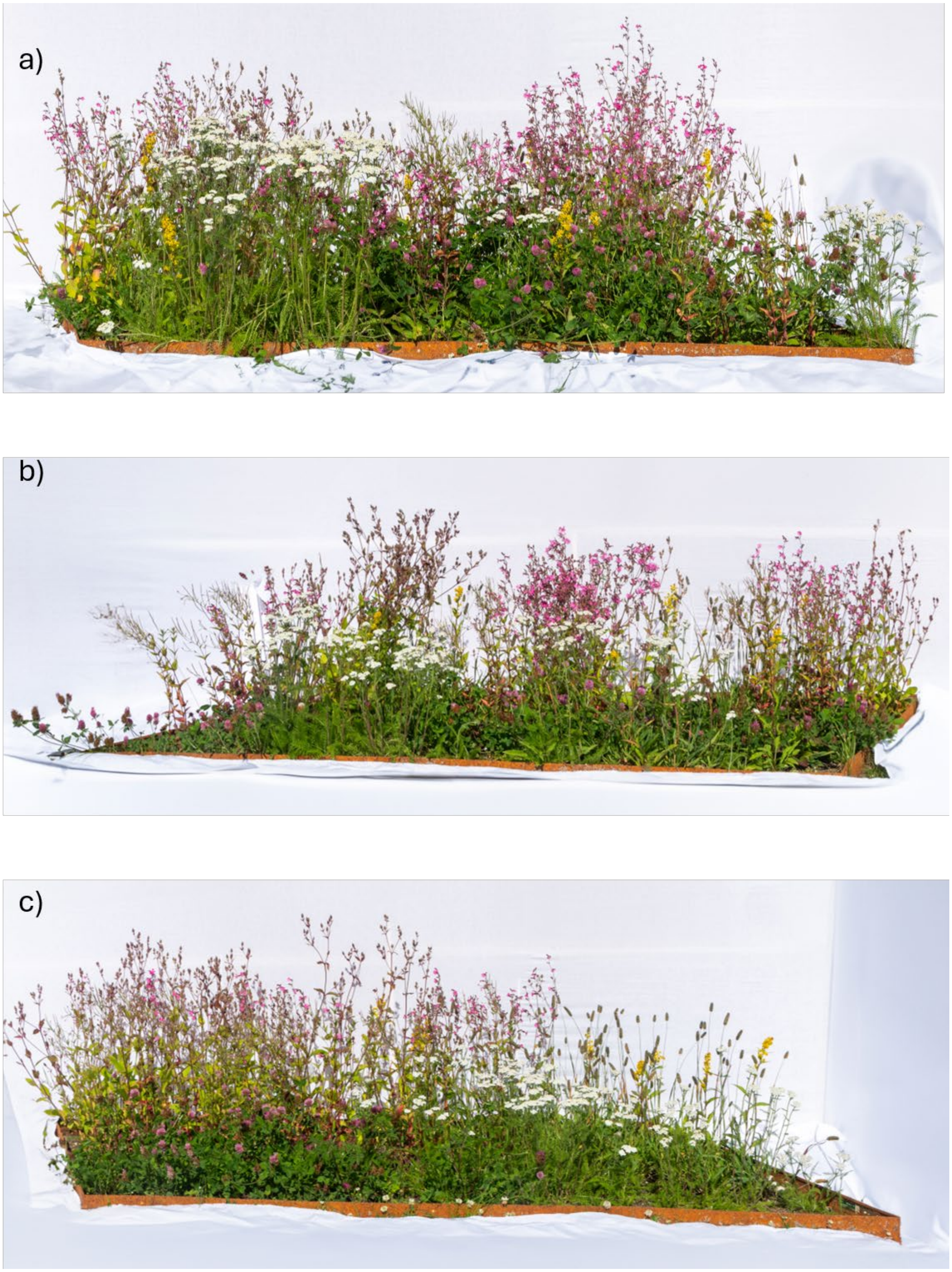


***Figure S2:*** *Flowerbeds in Set 1 (8 species) in no order (a), semi-order (b) and full order (c). Photo credit: Oddleif Larsen.*

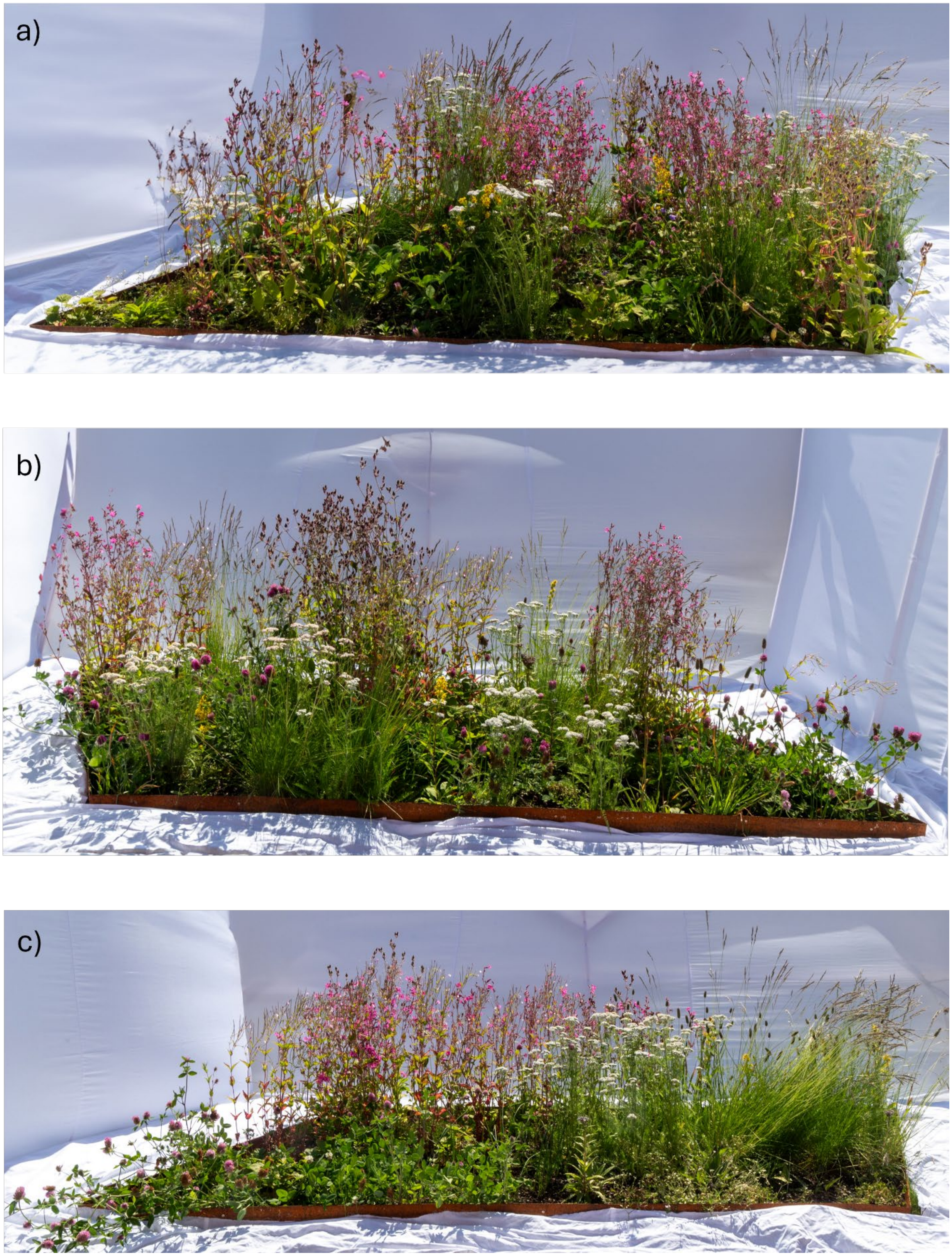


***Figure S2:*** *Flowerbeds in Set 3 (12 species) in no order (a), semi-order (b) and full order (c) Photo credit: Oddleif Larsen..*

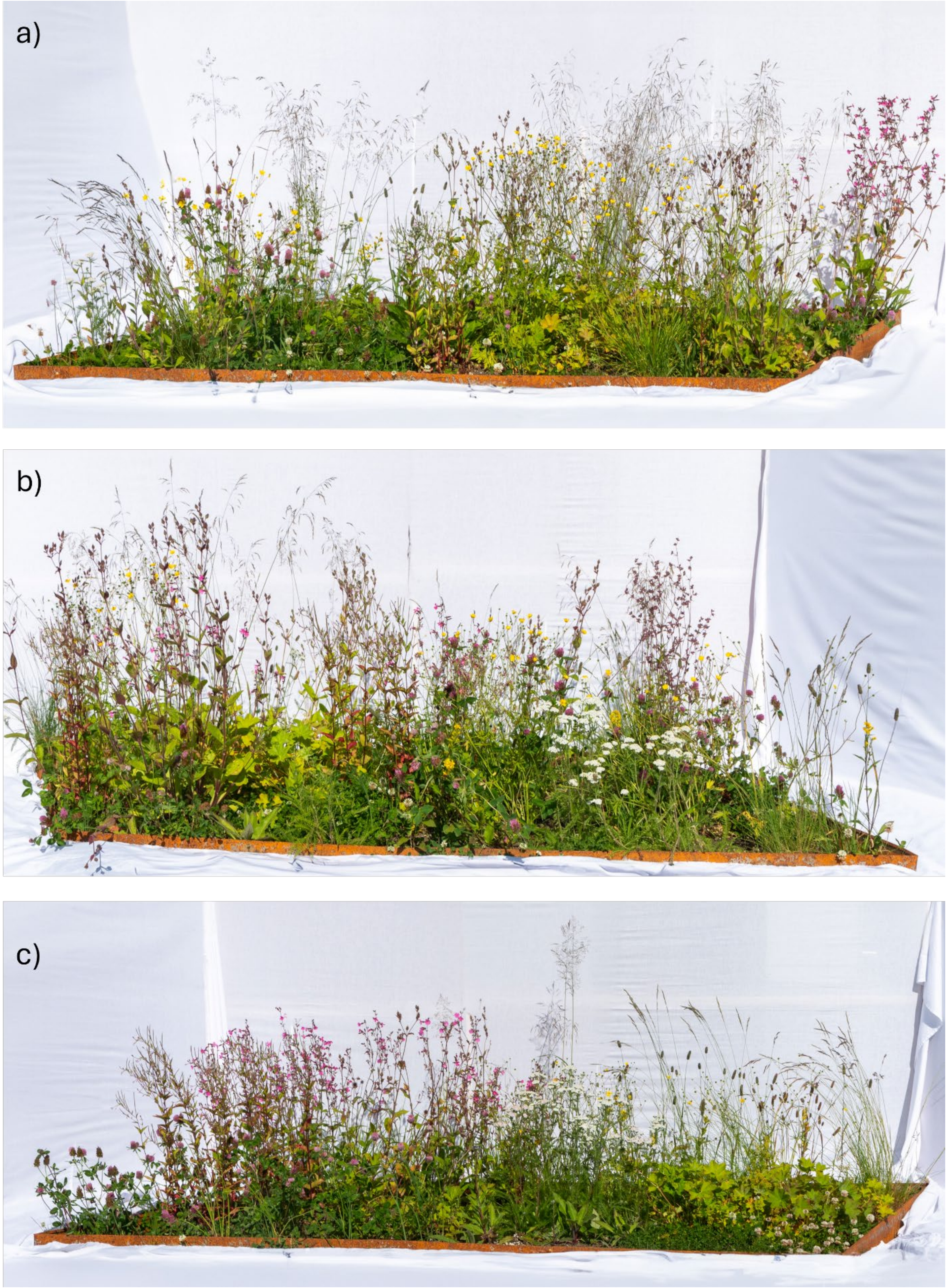


***Figure S3:*** *Flowerbeds in Set 2 (16 species) in no order (a), semi-order (b) and full order (c). Photo credit: Oddleif Larsen.*

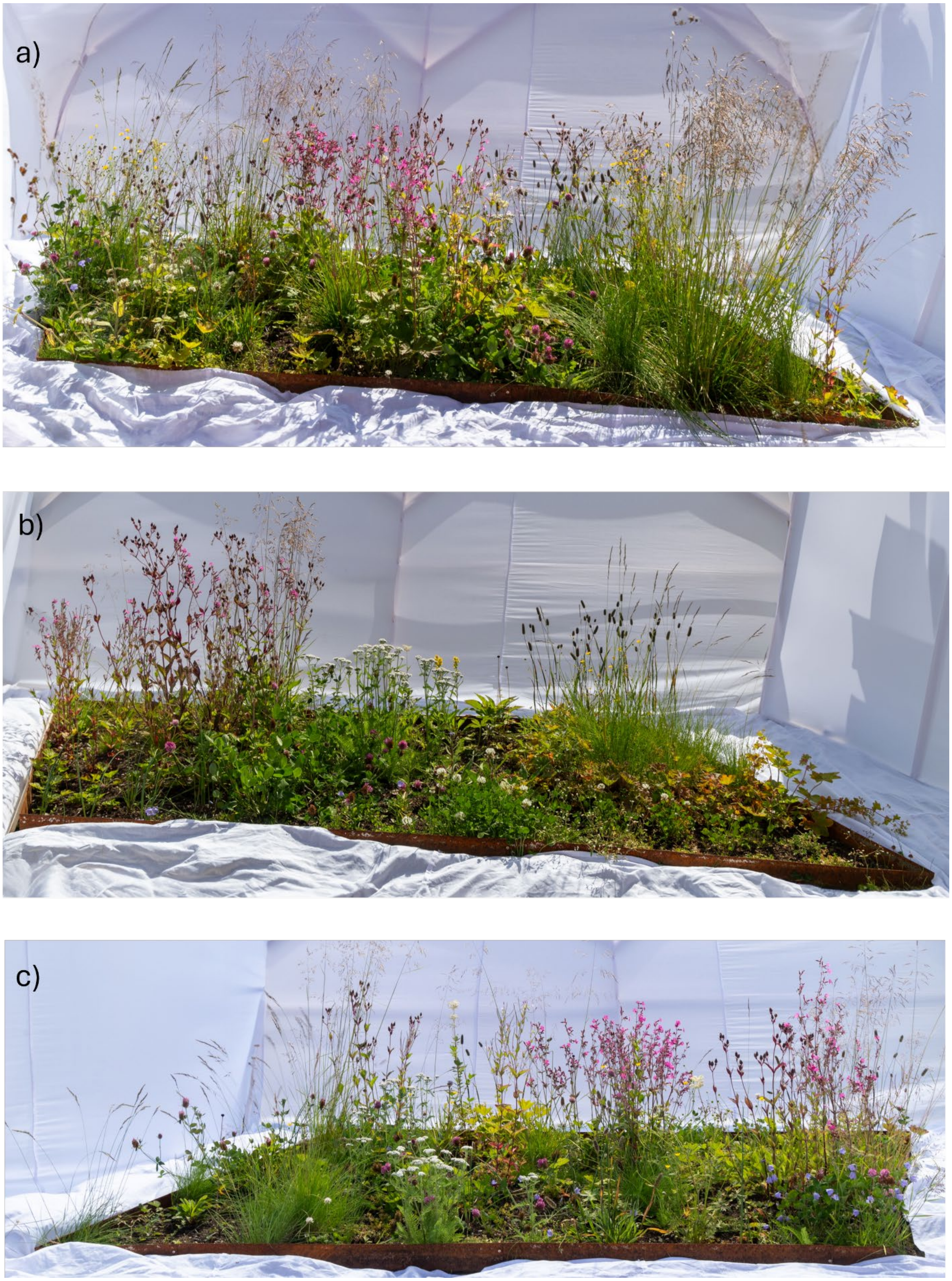


***Figure S4:*** *Flowerbeds in Set 4 (20 species) in no order (a), semi-order (b) and full order (c). Photo credit: Oddleif Larsen.*

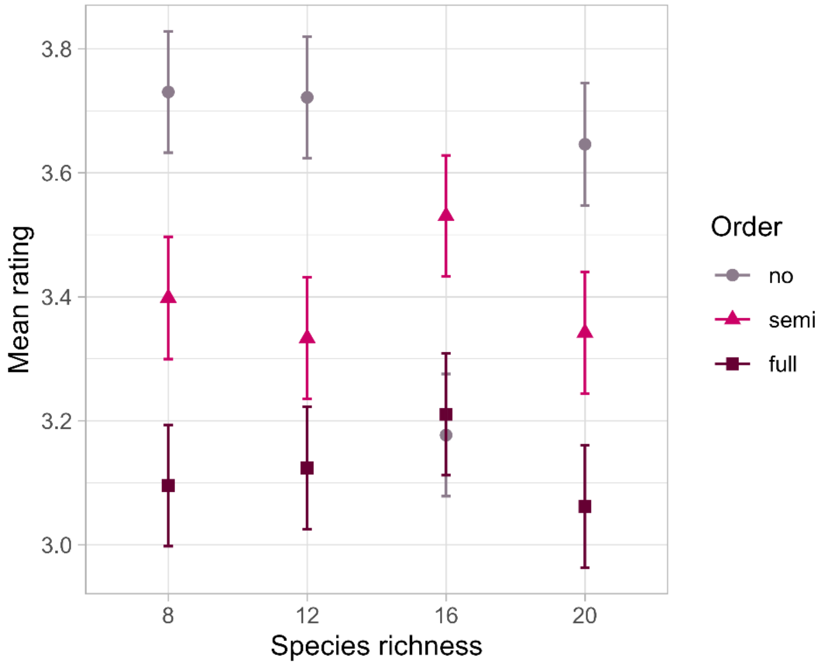


***Figure S5:*** *Average ratings (arithmetic mean ± standard error) for all plantings in the subset of answers* **excluding ecologists***. Colour and shape indicate the order of the planting*

***Table S3:*** *Model estimates for the subset of answers* ***excluding ecologists****. Generalized ordinal regression model (effect of species richness and order on ratings). The values given relate to an increasing order of ratings from negative to positive. Confidence intervals at the 95% level (CI). CIs not overlapping one indicate significant effects and are marked with **

| RATING increasing [negative/neutral/positive] | **P (Y=negative)** | **CI** $_{2.5}$ | **CI** $_{97.5}$ | **P (Y=negative/neutral)** | **CI** $_{2.5}$ | **CI** $_{97.5}$ |
|---|---|---|---|---|---|---|
| **INTERCEPT** | 0.1604* | 0.0992 | 0.2593 | 0.6016* | 0.4093 | 0.8841 |
| **SPECIES RICHNESS** | 1.0214 | 0.9920 | 1.0517 | 1.0104 | 0.9864 | 1.0350 |
| **ORDER – SEMI** | 0.9956 | 0.7089 | 1.3984 | 1.4004* | 1.0778 | 1.8196 |
| **ORDER – FULL** | 1.8112* | 1.3214 | 2.4826 | 2.4577* | 1.8824 | 3.2088 |

### Further findings

Additional to the findings based on the survey, we observed functional benefits of our plantings designed based on ecological principles. Growing seasons 2023 and 2024 were characterized by abnormally high temperatures (up to 28,2 °C in July 2023; 28,0 °C July 2024) and very low rainfall (18,4 mm in July 2023; and 4,7 mm in July 2024). This required watering of plantings on campus close to daily. We, however, refrained from watering the flowerbeds in 2024. The flowerbeds proved resilient against the severe drought, flowering lushly. The only negative consequence was an early onset of seed maturity and following senescence. In 2023 we offset this process with a mid-season cut of the seedbearing plants, which prompted their compensatory growth and re-production of flowers. The mid-season cut extended the flowering season even in severe drought conditions. In 2024 we omitted the midseason cut with a resulting shorter flowering season. We therefore recommend a midseason cut which can benefit appearance but also stability of the planting additional to the end-of-season cut as a measure of intermediate disturbance to avoid dominance of highly competitive species. Nevertheless, our plantings flowered earlier in spring and senescenced earlier in the fall than the surrounding plantings reliant on exotic species. We therefore suggest a selection of species that includes more late-flowering species to further elongate the flowering in the plantings and to include the appearance of the senescenced plant as an aesthetic factor in the plant selection process. The plants chosen for our plantings displayed mainly appealing fall appearances apart from T*rifolium pratense*, with dark brown leaves and stems in fall.
